# Recognition of β-D-glucose by surfactant protein D and structural evidence for an extended carbohydrate binding site

**DOI:** 10.64898/2026.11.24.768801

**Authors:** Jamie R. Littlejohn, Tania Castillo-Hernandez, Harry M. Williams, Amy J. Shaw, Yrina Ghrabigi, Georgina Brown, Howard W. Clark, Alastair Watson, Ian R. Burns, Kenneth B.M. Reid, Jens Madsen, Trevor J. Greenhough, Annette K. Shrive

## Abstract

Human surfactant protein D (hSP-D) is an innate immune protein involved in pathogen recognition in the lungs. Structural definition of the recognition of small carbohydrates by rfhSP-D, a biologically and therapeutically active recombinant homotrimeric fragment of hSP-D, has previously revealed the details of calcium dependent binding in the carbohydrate recognition domain and the influence of residues flanking the binding pocket.

To provide further structural insight into pathogen exo-polysaccharide recognition by hSP-D and the impact of alternative anomers and linkages, we have used x-ray crystallography and a competitive mannan-based ELISA to investigate the binding of α-D-glucose chains (3-7 sugars) and of β-linked glucose trisaccharides derived from long-chain cell wall components of fungal species and allergenic plant components and from extracellular matrices of bacterial biofilms.

Our results reveal that a variety of determinants influence the binding configuration, and that SP-D can recognise reducing terminal β-D-glucose through calcium mediated coordination of the O2ʹ and O1ʹ hydroxyls, with the flexibility to recognise alternative epitopes when the preferred target is unavailable for binding.

The ligands based on naturally occurring exo-polysaccharides reveal variable, novel and flexible recognition by hSP-D. β-cellotriose and β-laminaritriose exhibit different β-glucose binding modes (non-reducing O3ʹ, O4 ʹ and reducing O1ʹ, O2 ʹ respectively), both with moderate affinity for rfhSP-D. The high affinity β-isomaltotriose ligand binds via the reducing terminal β-glucose O2ʹ and O1ʹ in the calcium-binding pocket, supplemented by interaction of the internal, non-terminal glucose with an extended binding surface, revealing a potential mechanism for high affinity dual-site recognition of extended dextrans.

## Introduction

The collagenous C-type lectin surfactant protein D (SP-D) has long been associated with the innate immune response and activation of the acquired immune system (**1**), opsonising and clearing a wide range of pathogens through recognition of cell surface carbohydrate arrays (**2,3**). The localisation of hSP-D in the lung, circulatory system, saliva, and intestinal lumen, and its ability to regulate the immune response, places hSP-D in an ideal position to recognise these polysaccharides and contribute to the host response. Detailed investigation of the functional roles of SP-D is underpinned by three-dimensional structural studies of the biologically active recombinant homotrimeric fragment of human SP-D (rfhSP-D), comprising a short collagen-like domain, the α-helical coiled-coil neck plus three carbohydrate recognition domains (CRDs) (**4,5**). Structural definition of pathogen recognition has focused on small representative saccharide ligands, all bound via calcium and a terminal sugar hydroxyl pair in a mannose-type O3ʹ, O4ʹ configuration (**6,7**), and more recently on the recognition of gram-negative bacterial lipopolysaccharide (LPS), including *Haemophilus influenzae* (**8**) and *Salmonella enterica* Minnesota (**9**).

The recognition by SP-D of extended polysaccharides, often found in the cell walls or extra-cellular matrices of pathogenic or allergenic species such as plants, fungi and bacteria, has yet to be fully explored. Cellulose is a large, linear polymer of β(1–4)-linked glucoses found in the cell walls of most plants, some algae and in the connective material of a number of bacterial biofilms, including *Escherichia coli* and *Salmonella* strains (**10**). Class 1 α(1–6) dextrans are extracellular, glucose-rich polysaccharides that play an important role in bacterial cell adhesion and phagocyte evasion, contributing directly to the pathogenicity of *Streptococcus sp.* and the formation of dental plaques by *Streptococci* and *Lactobacilli* (**11**). The β(1–3)-linked β-glucan polymers are found in a wide range of organisms, most importantly in potentially pathogenic bacteria that affect humans (**12**). The roles of these β-glucans range from storage polysaccharides in protozoa and algae to cell wall components in fungi and yeast, including *Aspergillus fumigatus*. The recombinant hSP-D fragment rfhSP-D has been shown to recognise *A. fumigatus* cell surface β-glucan and to attenuate inflammation and offer protection in allergic asthma (**13**). In higher order plants the linear β-glucan callose forms an important part of the growth plates in the cell wall and along with cellulose is intrinsic to the cell wall of pollen tubes (**14**). A similar, largely linear form of β-glucan, curdlan, has been identified in the extracellular capsule of a number of medically important bacteria and has been implicated in the virulence of bacteria in other animals (**15,16**).

Recognition by rfhSP-D of the inner core LPS of a rough mutant of *H. influenzae* Eagan, 4A, occurs at the primary calcium site through heptose O6ʹ and O7ʹ hydroxyls alongside interaction of the anhydro 3-deoxy-D-manno-oct-2-ulosonic acid (Kdo) with the binding pocket flanking residues Arg343 and Asp325 (**8**). Structural definition of the binding of rfhSP-D to *S. enterica* Minnesota rough mutants R5 and R7 LPS (**9**) and to synthetic oligosaccharides (**17**) further revealed that SP-D has the versatility to adopt alternative recognition mechanisms when the inner core heptose is inaccessible. Binding of glycosylated structures on influenza virus has been investigated using molecular dynamics and structural studies, a hSP-D D325A+R343V mutant bound to a mannose-9-mer (**18**) showing recognition via Ca1 coordination of the O3ʹ, O4ʹ hydroxyls of the terminal mannose.

Several studies have reported variations and extensions beyond the standard recognition of a mannose-type O3ʹ, O4ʹ hydroxyl pair. Binding and structural studies of the recognition of α-D-glucosyl trisaccharides demonstrated an extended binding surface beyond the calcium site involving Thr336 and Asn337 along with a π-stacking interaction between the aromatic ring of Phe335 and the third, terminal moiety of the three membered chain (**19**). This is reflected in the increased affinity of SP-D for β-maltotriose and *p*-nitrophenyl-α-D-maltoside (and to a lesser extent β-maltotetraose) compared to maltose (**19**). Binding of non-calcium bound terminal sugars has also been demonstrated for a rfhSP-D R343V mutant complexed with α1-2 dimannose (**6**) where the non-calcium bound mannose interacts at a secondary site including Glu347 and Arg349. In terms of non-calcium bound non-terminal sugars, based on the structures of rfhSP-D complexed with the inner core oligosaccharide of the *S. enterica* sv Minnesota rough strains R5 and R7 (**9**), an extended ligand binding site, in the region of the SP-D conserved Glu333, Glu347 and Arg349, has been proposed. Molecular modelling studies have suggested that SP-D recognition of glucosyl trisaccharides in the calcium-binding pocket can also occur via non-terminal glucosyl residues (**20**). Internal glucosyl residue binding was proposed for β-isomaltotriose (α1-6) and β-gentiotriose (β1-6) via the 2ʹ, 3ʹ and 3ʹ, 4ʹ hydroxyl pairs respectively and for β-maltotriose (α1-4 linked, 2ʹ, 3ʹ hydroxyls), but not for β-cellotriose (β1-4) or for 1-3 linked glucosyl residues in general (**20**).

We have investigated the interaction of the longer α-D-glucose chains β-maltotetraose and β-maltoheptaose and of physiologically relevant oligosaccharides derived from long-chain cell wall components of fungal species and allergenic plant components and from extracellular matrices of bacterial biofilms (**11**). hSP-D binding of ligands derived from cellulose and β-glucan, β-cellotriose (β(1–4)-linked) and β-laminaritriose (β(1–3)-linked) respectively, has been characterised along with β-isomaltotriose (α(1–6)-linked) from dextran, to provide further structural insight into pathogen exo-polysaccharide recognition and the impact of alternative anomers and linkages, and of non-terminal sugars (**20**), on carbohydrate binding by hSP-D.

## Results

The structures of the carbohydrate ligands studied here along with selected relevant ligands from previous studies are shown in Table **1**.

**Table 1.** Names and structures for each carbohydrate ligand studied here, along with selected relevant ligands from previous studies, in the chair pyranose structure. Glycosidic bonds and configurations are shown in red, alongside the principal anomeric form of the C1 carbon. Figures produced using ChemDraw.

| Carbohydrate | Chair Pyranose Structure |
|---|---|
| β-Maltotriose<br>α-D-Glc-(1,4)-α-D-Glc-(1,4)-β-D-Glc |  |
| β-Maltotetraose<br>[α-D-Glc-(1,4)]<sub>3</sub>-β-D-Glc |  |
| β-Maltoheptaose<br>[α-D-Glc-(1,4)]<sub>6</sub>-β-D-Glc |  |
| β-Isomaltotriose<br>α-D-Glc-(1,6)-α-D-Glc-(1,6)-β-D-Glc |  |
| β-Laminaritriose<br>β-D-Glc-(1,3)-β-D-Glc-(1,3)-β-D-Glc |  |
| β-Cellotriose<br>β-D-Glc-(1,4)-β-D-Glc-(1,4)-β-D-Glc |  |
| β-Galactose<br>β-D-Gal |  |
| N-acetyl-α-D-mannosamine<br>(ManNAc) |  |
| Maltose<br>α-D-Glc-(1,4)-α-D-Glc |  |

### Binding studies

The binding affinity of rfhSP-D to different carbohydrates in presence of calcium was assessed using a competitive mannan-based enzyme linked immunosorbent assay (ELISA). The binding between rfhSP-D and mannan was inhibited by the presence of different carbohydrates in a concentration-dependent manner (Fig. **1A**, Supplementary Fig. **S1**). This binding was completely inhibited by the presence of 10 mM EDTA in solution or 12.5 mM maltose and 6.5 mM ManNAc (Supplementary Fig. **S2**). The relative half-maximal inhibitory concentrations of each carbohydrate (IC_50_) are summarised in Fig. **1B** and Table **2**. The most effective inhibitors were maltotriose, ManNAc, and isomaltotriose followed by cellotriose, maltoheptaose, maltose and laminaritriose with lesser degree of inhibition. Finally, galactose had the least inhibitory effect of all tested carbohydrates. The results show relative binding affinity for rfhSP-D as: Maltotriose > ManNAc > Isomaltotriose > Cellotriose > Maltoheptaose > Maltose > Laminaritriose >> Galactose.

**Figure 1.**
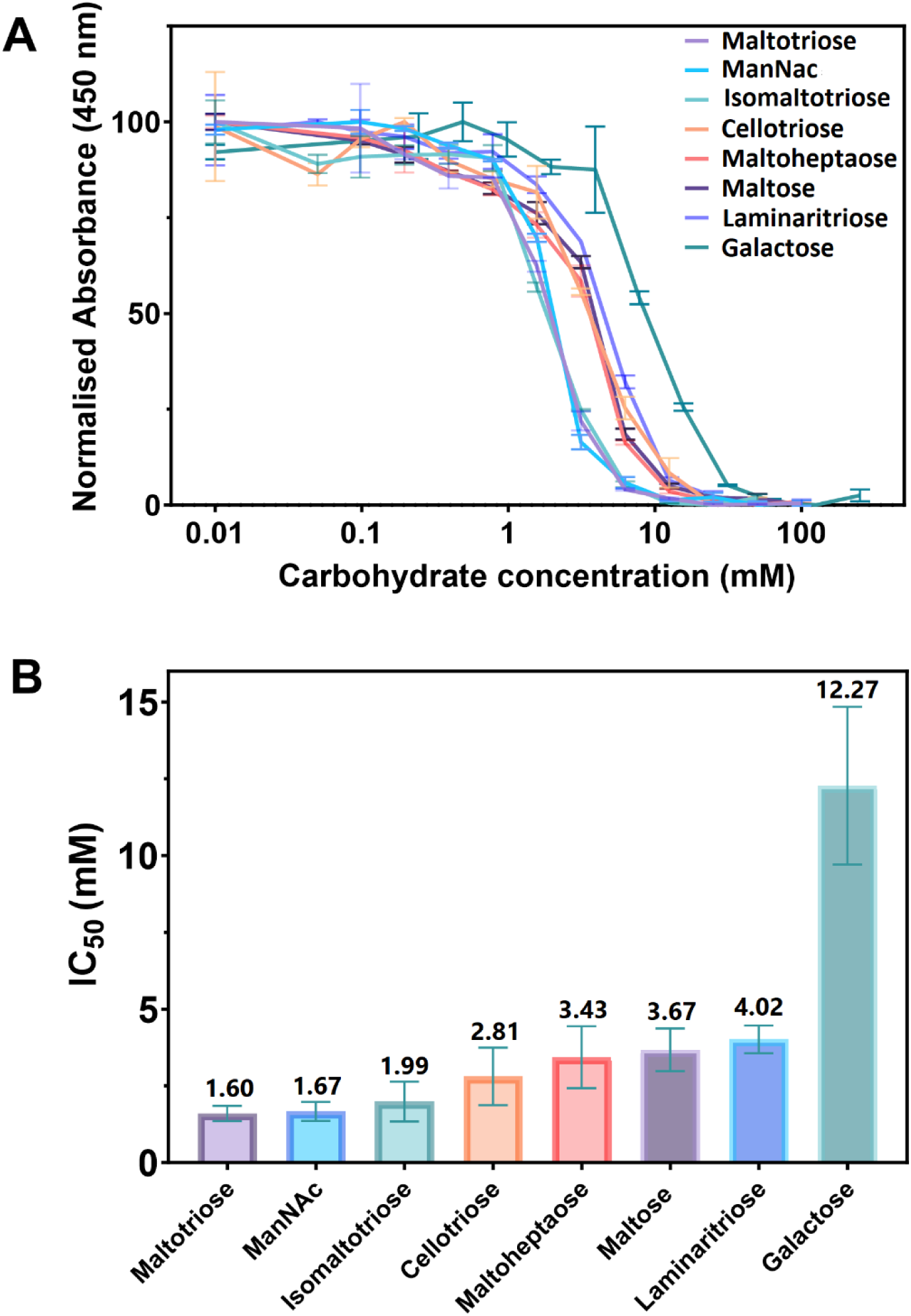
Binding affinities of rfhSP-D to different carbohydrates. (**A**) The IC_50_ values represent the average concentration of each carbohydrate required to inhibit 50% of the binding of rfhSP-D to mannan. (**B**) IC_50_ values obtained using a competitive mannan-based ELISA. Data shown as mean ± standard deviation of 3-6 independent experiments.

**Table 2.**
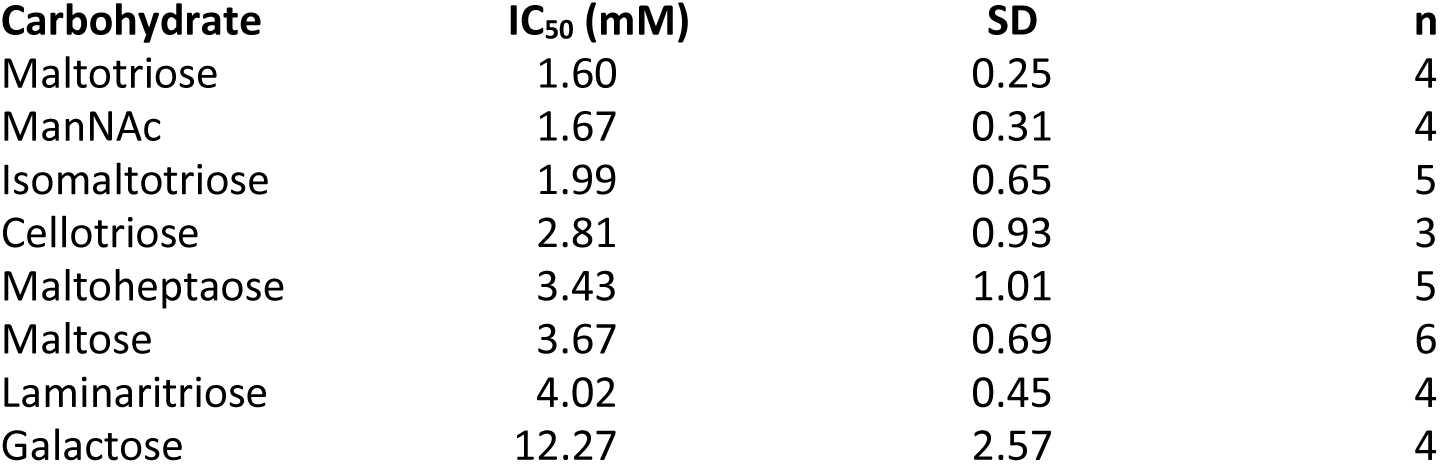
IC_50_ values for the binding of rfhSP-D to different carbohydrates. IC_50_ values obtained using a competitive mannan-based ELISA. Data shown as mean ± standard deviation of n independent experiments.

### Crystal structures

The high-resolution crystal structures of the trimeric neck-CRD hSP-D fragment rfhSP-D bound to β-maltotetraose, β-maltoheptaose, β-cellotriose, β-isomaltotriose and β-laminaritriose have been determined. The overall protein structures are similar to those published previously, with the ligand binding site in all three subunits A, B and C accessible to ligand but more restricted in subunit A due to the close approach of a neighbouring subunit in the crystal (**21**). Protein coordination to both the primary calcium Ca1 (via Glu321, Asn323, Glu329, Asn341) and the calcium-bound ligand hydroxyls (via Glu321, Asn323, Glu329, Asn341, Asp342) follows the established pattern for ligand-bound SP-D structures (Tables **3, S1**). The binding-pocket flanking residue Asp325 is in a similar position and orientation in all structures with some heterogeneity in the orientation of the carboxyl head group. Ligand structures and their interactions with rfhSP-D are summarised in Table **4**. A fourth calcium ion, Ca4, located on the trimeric axis at the bottom of the funnel formed by the three CRDs is present in the β-maltotetraose and β-maltoheptaose bound structures due to addition of calcium into the cryobuffers (**21**).

### β -maltoheptaose

The binding of β-maltoheptaose to SP-D is calcium mediated via the equatorial hydroxyl groups O3ʹ and O4ʹ of the non-reducing terminal α-D-glucose Glc1 in subunit A (2.48, 2.59 Å) and O2ʹ, O1ʹ of the reducing terminal β-D-glucose (Glc7) in subunits B and C (2.53-2.61 Å). In subunit A the electron density shows the two terminal glucose residues Glc1 and Glc2, positioned as for maltose, both structures having two conformations of Arg343 in subunit A (**21**) (Fig. **2A**, **2B**). Glucose residues Glc3-Glc7 are not visible in the electron density. Subunits B and C show a single β-D-glucose (Glc7) (Fig. 2C) with the Glc7 pyranose ring rotated by 180° compared to the Glc1 in subunit A such that the pyranose face of the bound glucose presented to Asp325 in subunit A differs from that in subunits B and C. Despite this, O6ʹ of the bound glucose forms an interaction with Arg343 NH1 in all three subunits, at 3.18 Å, 3.27 Å and 3.25 Å in subunits A, B and C respectively (two conformations of Arg343 are present in subunit A – closest approach given).

**Figure 2.**
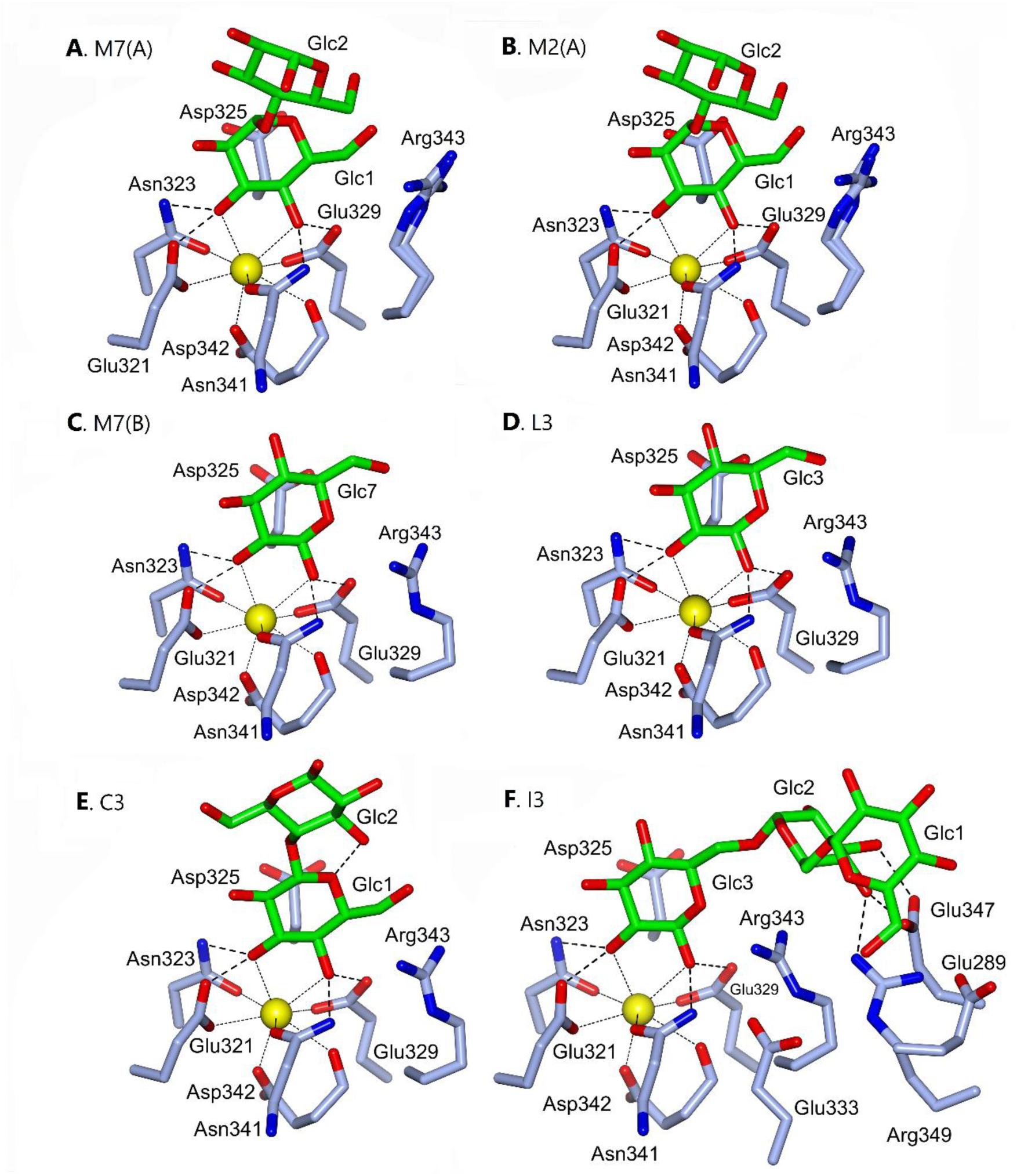
Binding of α-D-glucose chains and β-linked glucose trisaccharides to rfhSP-D. β-maltotetraose is shown in figure 3.**(A)** β-maltoheptaose (M7) and **(B)** maltose (M2, pdb 1pwb) bound in subunit A of rfhSP-D. Maltoheptaose Glc3-7 are not defined in the electron density. **(C)** β-maltoheptaose (M7) bound in subunit B of rfhSP-D, also representative of subunit C. Maltoheptaose Glc1-6 are not defined in the electron density. **(D)** β-laminaritriose (L3) bound in rfhSP-D subunit B. Glc1-2 are not defined in the electron density. Representative of all three subunits, with some heterogeneity in the position of O6ʹ. **(E)** β-cellotriose (C3) bound in rfhSP-D subunit A. Glc3 is not defined in the electron density with only partial density for Glc2 in subunits B and C. **(F)** β-isomaltotriose (I3) bound in rfhSP-D subunit A. Representative of all 3 subunits.

### β-laminaritriose

In all three subunits of the β-laminaritriose bound structure, only the reducing terminal β-D-glucose (Glc3) is observed, coordinating Ca1 through the O2ʹ and O1ʹ hydroxyls (Fig. **2D**). The orientation of Glc3 in the binding site positions Glc3 O6ʹ in proximity to Arg343 NH1 in subunit B (3.08 Å) with significantly longer distances in all three subunits between Arg343 and Glc3 O5. In subunits A and C O6ʹ is directed away from Arg343.

### β-cellotriose

In subunit A of the β-cellotriose (β(1–4)-linked) bound structure (Fig. **2E**) two of the three glucose rings are visible in the electron density, with the O3ʹ and O4ʹ hydroxyl groups of βGlc1 coordinating to Ca1 at distances of 2.45-2.61 Å and to protein in the established pattern for ligand-bound SP-D structures (Tables **3, S1**). Partial density for Glc2 in subunits B and C indicates that this sugar is similarly placed to that in subunit A. The pyranose face of the bound glucose presented to Asp325 is the opposite to that for β-laminaritriose, β-isomaltotriose and the B, C subunits of β-maltoheptaose, which all coordinate to calcium via the reducing terminal Glc equatorial O2ʹ and O1ʹ hydroxyls. The Glc1 O6ʹ-Arg343 NH1 distances are 2.61-3.29 Å across the three subunits.

### β-isomaltotriose

In all subunits the binding of β-isomaltotriose is via the reducing terminal Glc3 equatorial O2ʹ and O1ʹ hydroxyls (2.49-2.61 Å) with O5 and O6ʹ of Glc3 interacting with Arg343 at 2.96-3.32 Å and 3.41-3.46 Å respectively. This O2ʹ and O1ʹ hydroxyl coordination results in the α(1–6) linkage extending away from the binding site above Arg343, with Glc2 completing the protein-ligand interactions at a secondary site where Glc2 is recognised by Glu347 (2.46-3.01 Å) and Arg349 (2.66-3.03 Å) (Fig. **2F**) supplemented by a series of water bridges. In all three subunits there is density sufficient to define the orientation of Glc1 extending beyond the anchored internal Glc2, but the position of Glc1 O6ʹ and its interactions with protein cannot be modelled with confidence.

### β-maltotetraose

Recognition of β-maltotetraose is via the O3ʹ and O4ʹ hydroxyls of the non-reducing terminal glucose (Glc1) in all three subunits (Fig. **3**; Table **3**). Glc1 O6ʹ-Arg343 NH1 is 3.18-3.34 Å across the three subunits. In subunit A all four glucoses are seen with Glc2-4 extending away from the binding pocket, forming an arch across the protein surface with the reducing terminal β-glucose Glc4 anchored via Thr336 (O1ʹ Glc4-OG1 Thr336 2.87 Å) and the Asn337 main-chain N (O1ʹGlc4–NAsn337 2.88 Å) in a similar manner to the reducing terminal Glc in the maltotriose-bound structure (**19**). This is complemented by a water bridge between Asn337 and Glc4 and a water mediated H bond network between Glu321, Glu333, Arg343 and Glc2, which interacts with Glc3. There is little resolution of Glc2-Glc4 in subunits B and C beyond the glycosidic linkage extending from Glc1 O1ʹ.

**Figure 3.**
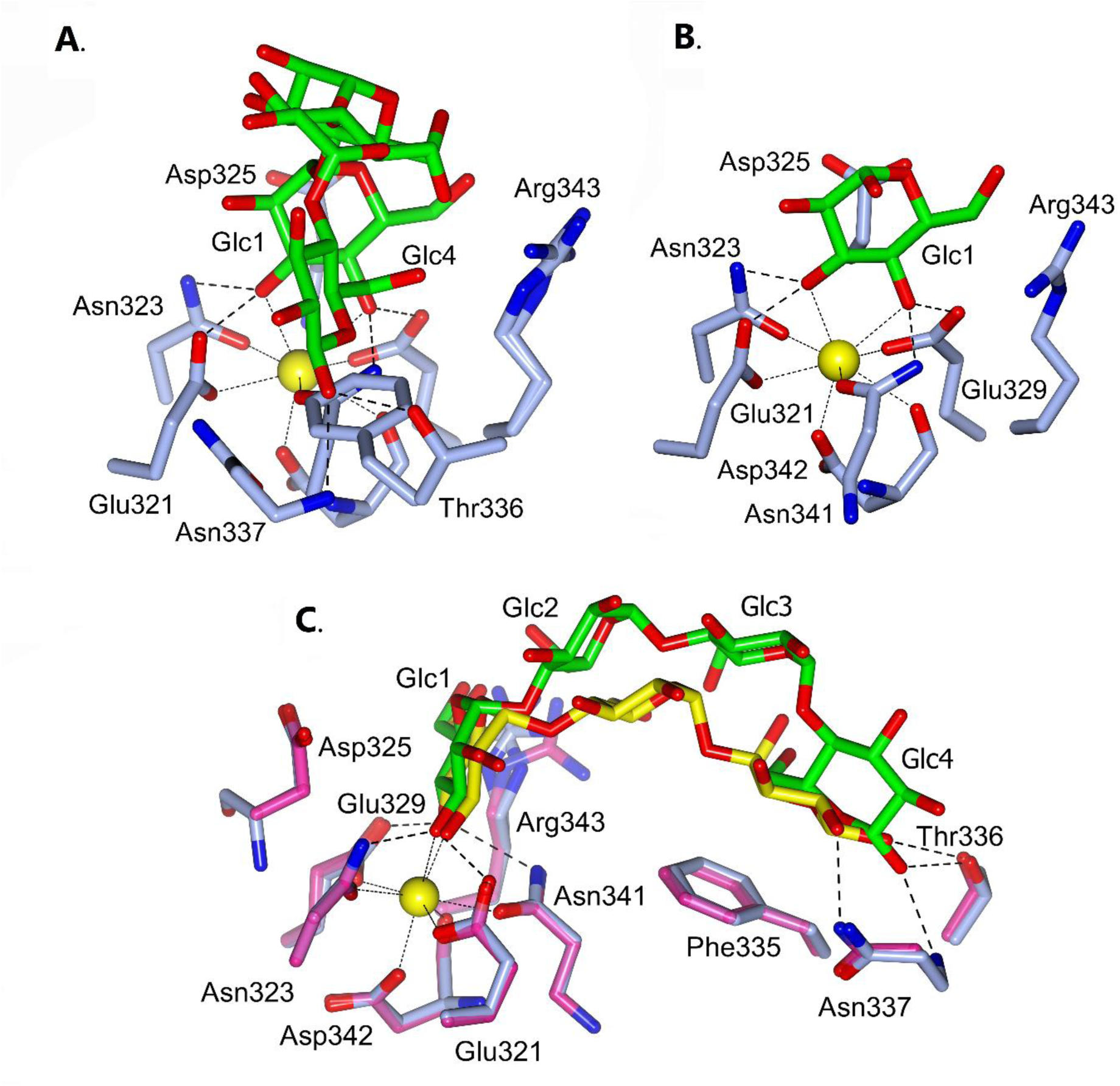
Binding of β-maltotetraose to rfhSP-D. β-maltotetraose bound in **(A)** rfhSP-D subunit A, **(B)** rfhSP-D subunit B. Glc2-4 are not defined in the electron density in subunit B. Subunit C is similar to subunit B. **(C)**: Subunit A orthogonal view showing bound maltotriose (pdb 2ggu, yellow) and maltotetraose (green). β-maltotriose-bound rfhSP-D protein is shown in Purple; β-maltotetraose in Blue.

**Table 3.** Calcium-ligand and protein-ligand binding distances (Å)

| **Atom 1** |  | **Atom 2** |  |  | **Maltotetraose (M4)** |  |  | **Maltoheptaose (M7)** |  |  | **Isomaltotriose (I3)** |  |  | **Cellotriose (C3)** |  |  | **Laminaritriose (L3)** |  |  |
|---|---|---|---|---|---|---|---|---|---|---|---|---|---|---|---|---|---|---|---|
|  |  |  |  |  | A | B | C | A | B | C | A | B | C | A | B | C | A | B | C |
| Ca1 |  | Glc |  | O3′ | 2.49 | 2.53 | 2.51 | 2.48 | - | - | - | - | - | 2.49 | 2.45 | 2.52 | - | - | - |
|  |  |  |  | O4′ | 2.63 | 2.59 | 2.63 | 2.59 | - | - | - | - | - | 2.54 | 2.61 | 2.57 | - | - | - |
|  |  | Glc |  | O1′ | - | - | - | - | - | - | 2.61 | 2.56 | 2.49 | - | - | - | 2.57 | 2.61 | 2.61 |
|  |  |  |  | O2′ | - | - | - | - | - | - | 2.52 | 2.49 | 2.54 | - | - | - | 2.54 | 2.48 | 2.58 |
|  |  | Glc |  | O1′ | - | - | - | - | 2.61 | 2.58 | - | - | - | - | - | - | - | - | - |
|  |  |  |  | O2′ | - | - | - | - | 2.53 | 2.53 | - | - | - | - | - | - | - | - | - |
| Glc1 | O3′ | Glu | 321 | OE2 | 2.61 | 2.51 | 2.55 | 2.59 | - | - | - | - | - | 2.46 | 2.55 | 2.54 | - | - | - |
|  |  | Asn | 323 | ND2 | 2.72 | 2.87 | 2.85 | 2.77 | - | - | - | - | - | 2.74 | 2.88 | 2.84 | - | - | - |
|  | O4′ | Glu | 329 | OE2 | 2.54 | 2.47 | 2.51 | 2.55 | - | - | - | - | - | 2.53 | 2.51 | 2.50 | - | - | - |
|  |  | Asn | 341 | ND2 | 2.88 | 3.03 | 2.95 | 2.88 | - | - | - | - | - | 3.00 | 3.02 | 3.01 | - | - | - |
| Glc3 | O1′ | Glu | 329 | OE2 | - | - | - | - | - | - | 2.62 | 2.47 | 2.56 | - | - | - | 2.55 | 2.58 | 2.58 |
|  |  | Asn | 341 | ND2 | - | - | - | - | - | - | 3.00 | 3.02 | 2.95 | - | - | - | 3.07 | 3.05 | 3.05 |
|  | O2′ | Glu | 321 | OE2 | - | - | - | - | - | - | 2.62 | 2.52 | 2.63 | - | - | - | 2.61 | 2.50 | 2.46 |
|  |  | Asn | 323 | ND2 | - | - | - | - | - | - | 2.83 | 2.81 | 2.87 | - | - | - | 2.88 | 2.91 | 2.89 |
| Glc7 | O1′ | Glu | 329 | OE2 | - | - | - | - | 2.52 | 2.51 | - | - | - | - | - | - | - | - | - |
|  |  | Asn | 341 | ND2 | - | - | - | - | 3.03 | 3.00 | - | - | - | - | - | - | - | - | - |
|  | O2′ | Glu | 321 | OE2 | - | - | - | - | 2.55 | 2.55 | - | - | - | - | - | - | - | - | - |
|  |  | Asn | 323 | ND2 | - | - | - | - | 2.88 | 2.84 | - | - | - | - | - | - | - | - | - |
| Glc4 | O1′ | Thr | 336 | OG1 | 2.87 | - | - | - | - | - | - | - | - | - | - | - | - | - | - |
|  |  | Asn | 337 | N | 2.88 | - | - | - | - | - | - | - | - | - | - | - | - | - | - |
| Glc2 | O3′ | Glu | 347 | OE1 | - | - | - | - | - | - | 3.01 | - | - | - | - | - | - | - | - |
|  |  | Glu | 347 | OE2 | - | - | - | - | - | - | 2.82 | 2.50 | 2.46 | - | - | - | - | - | - |
|  | O4′ | Glu | 347 | OE1 | - | - | - | - | - | - | 2.47 | 2.96 | - | - | - | - | - | - | - |
|  |  | Arg | 349 | NH2 | - | - | - | - | - | - | 2.66 | 3.03 | 2.96 | - | - | - | - | - | - |
| Glc3 | O5 | Arg | 343 | NH1 | - | - | - | - | - | - | 3.08 | 3.06 | 2.96 | - | - | - | - | - | - |
| Glc3 | O5 | Arg | 343 | NH2 | - | - | - | - | - | - | 3.24 | 3.32 | 3.23 | - | - | - | - | - | - |
| Glc1 | O6′ | Arg | 343 | NH1 | 3.20 | 3.34 | 3.34 | 3.18 | - | - | - | - | - | 2.61 | 3.29 | 2.86 | - | - | - |
| Glc3 | O6′ | Arg | 343 | NH1 | - | - | - | - | - | - | 3.41 | 3.46 | 3.45 | - | - | - | 3.26 | 3.08 | 3.41 |
| Glc7 | O6′ | Arg | 343 | NH1 | - | - | - | - | 3.27 | 3.25 | - | - | - | - | - | - | - | - | - |

<sup>1</sup> Arg343 has two conformations in subunit A of M4 and M7. Closest approach shown

**Table 4.** Summary of ligands and interactions.

| Ligand | Main component | Linkage | Number of glucose residues | Reducing terminal anomer | Ca2+ recognition residue |  |  | Ca2+ hydroxyl coordination |  |  | Number of defined glucose residues |  |  | Extended binding site |  |  | Overall Chain Structure |
|---|---|---|---|---|---|---|---|---|---|---|---|---|---|---|---|---|---|
|  |  |  |  |  | Subunit |  |  | Subunit |  |  | Subunit |  |  | Subunit |  |  |  |
|  |  |  |  |  | A | B | C | A | B | C | A | B | C | A | B | C |  |
| β-cellotriose | β-glucose | β(1→4) | 3 | β | β Glc1 | β Glc1 | β Glc1 | O3' O4' | O3' O4' | O3' O4' | 2 | 1 | 1 | None | None | None | [β-D-Glc-(1→4)]<sub>2</sub>-β-D-Glc |
| β-isomaltotriose | α-glucose | α(1→6) | 3 | β | β Glc3 | β Glc3 | β Glc3 | O2' O1' | O2' O1' | O2' O1' | 2* | 2* | 2* | αGlc2 Glu347 Arg349 | αGlc2 Glu347 Arg349 | αGlc2 Glu347 Arg349 | [α-D-Glc-(1→6)]<sub>2</sub>-β-D-Glc |
| β-laminaritriose | β-glucose | β(1→3) | 3 | β | β Glc3 | β Glc3 | β Glc3 | O2' O1' | O2' O1' | O2' O1' | 1 | 1 | 1 | None | None | None | [β-D-Glc-(1→3)]<sub>2</sub>-β-D-Glc |
| maltose<br>pdb 1pwb | α-glucose | α(1→4) | 2 | α | α Glc1 | α Glc1 | α Glc1 | O3' O4' | O3' O4' | O3' O4' | 2 | 1 | 1 | None | None | None | α-D-Glc-(1→4)-α-D-Glc |
| β-maltotriose<br>pdb 2ggu | α-glucose | α(1→4) | 3 | β | α Glc1 | α Glc1 | α Glc1 | O3' O4' | O3' O4' | O3' O4' | 3 | 3 | 3 | βGlc3 Phe335 Thr336 Asn337 | βGlc3 Phe335 Thr336 Asn337 | βGlc3 Phe335 Thr336 Asn337 | [α-D-Glc(1→4)]<sub>2</sub>-β-D-Glc |
| β-maltotetraose | α-glucose | α(1→4) | 4 | β | α Glc1 | α Glc1 | α Glc1 | O3' O4' | O3' O4' | O3' O4' | 4 | 1 | 1 | βGlc4 Thr336 Asn337 | None | None | [α-D-Glc-(1→4)]<sub>3</sub>-β-D-Glc |
| β-maltoheptaose | α-glucose | α(1→4) | 7 | β | α Glc1 | β Glc7 | β Glc7 | O3' O4' | O2' O1' | O2' O1' | 2 | 1 | 1 | None | None | None | [α-D-Glc-(1→4)]<sub>6</sub>-β-D-Glc |

\* Glc1 has poor overall definition in the electron density, but the orientation and position of the Glc1 ring can be assigned.

## Discussion

The series of structures in this study reveals important mechanisms for carbohydrate ligand recognition by surfactant protein D, providing an insight into the recognition of naturally occurring polysaccharides, particularly those which include alternative glycosidic linkages. We have previously shown for bacterially derived LPS that SP-D is able to recognise internal inner core sugars at the calcium site and to utilise both an extended binding site and alternative epitopes of the same ligand when the preferred target is not available for binding (**9**). The β-**maltoheptaose** (α1-4 linked) structure (Fig. **2A****, 2C**) presented here demonstrates alternative modes of binding while the α1-6 linked β-**isomaltotriose** (Fig. **2F**) utilises an extended binding surface for non-calcium coordinated internal sugars which may enhance recognition of longer oligosaccharide ligands containing alternative linkages. β**-laminaritriose** (β1-3 linked) and β**-cellotriose** (β1-4 linked), the first poly β-D-glucose ligands to be investigated bound to rfhSP-D, adopt different binding modes with β**-laminaritriose** bound via βGlc3 O2ʹ and O1ʹ (reducing) and β**-cellotriose** via βGlc1 O3ʹ and O4ʹ. For ligands where binding studies have been carried out previously, the binding data presented here (Fig. **1****, S1, S2)** broadly agree overall with previously published data (**19,22**) where high affinity binding to maltotriose and ManNAc was reported.

Overall, the configuration of the glucose bound at the calcium site, and of Arg343 (**6,9,21**), fall into 3 distinct groups (Fig. **4**) categorised by the pyranose face presented to Asp325 (**23,24**), the pair of hydroxyls bound and the nature of the linkage to the next sugar. In groups A and B (as defined in Fig. **4**) calcium coordination is via the non-reducing terminal O3ʹ, O4ʹ while in group C coordination is via the reducing terminal O2ʹ, O1ʹ with the opposite pyranose face to groups A and B presented to Asp325. In addition, the nature of the linkage to the next sugar is α in group A, β in group B, and both in group C. The configurations of the calcium and the glucose coordinating residues, and of the flanking residue Asp325, are closely similar across all the subunits and structures with the conformation of Arg343 in groups B and C differing from that in group A where subunit A of **maltose,** β**-maltotetraose** and β-**maltoheptaose (group A)** show dual Arg343 conformations.

**Figure 4.**
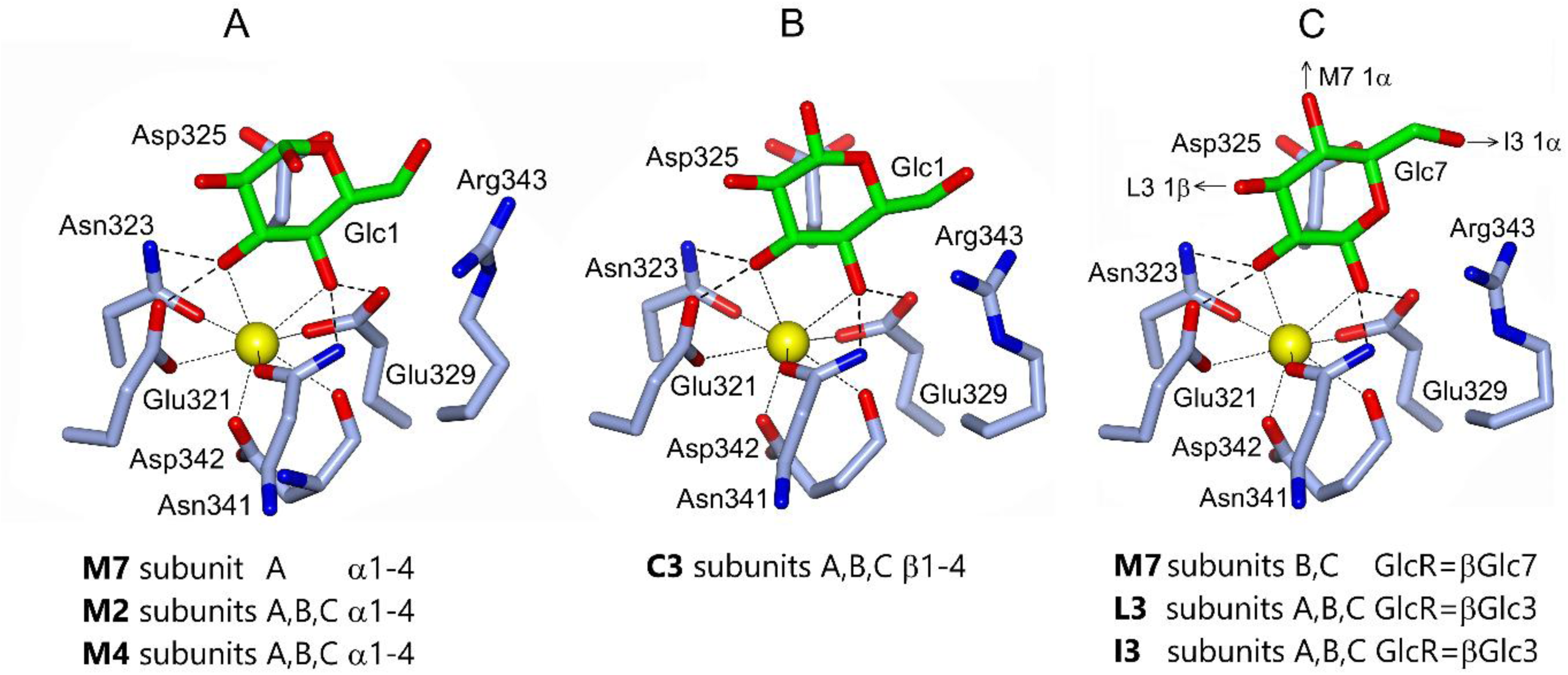
The three glucose binding modes at the calcium binding site. Structures shown are representative of the ligand-bound structures in each group. Only the terminal, bound pyranose is shown. **(A)** the non-reducing terminal α glucose of β maltoheptaose (M7), maltose (M2, pdb 1pwb) and β -maltotetraose (M4). In each case the dual conformation of Arg343 in subunit A is not shown for clarity. **(B)** the non-reducing terminal β glucose of β-cellotriose (C3) **(C)** the reducing terminal β glucose of β-maltoheptaose (M7), β-laminaritriose (L3) and β-isomaltotriose (I3).

The O2ʹ and O1ʹ hydroxyls of the reducing terminal β-D-glucose Glc3 in all three subunits of β-**isomaltotriose** (α(1–6)-linked) and β-**laminaritriose** (β(1–3)-linked), and of β-**maltoheptaose** Glc7 (α(1–4)-linked) in subunits B and C, are bound by rfhSP-D in a novel β-D-glucose mediated mechanism (Fig. **4C**) which is similar to the binding of **galactose** by the same rfhSP-D (**7**). These three structures suggest selectivity for O2ʹ and O1ʹ hydroxyls when they are available since for both β-**laminaritriose** and β-**isomaltotriose** the Glc1 O3ʹ and O4ʹ hydroxyls are available on the non-reducing terminal glucose residue (and for β-**isomaltotriose** also on the reducing terminal β-glucose) but do not coordinate the calcium in this way as do β-**maltotetraose**, **maltose,** β-**cellotriose** and β-**maltotriose** (Fig. **4A**, **4B**) (**19,21**). This is particularly evident in the β-**maltoheptaose**-bound structure where both binding mechanisms are present in the crystal structure (Fig. **2A**, **2C**). Recognition of O2ʹ, O1ʹ (reducing terminal β-glucose Glc7) in the more accessible and open β-**maltoheptaose** subunits B and C sites again suggests this binding is preferred to the non-reducing terminal O3ʹ, O4ʹ seen in the more constricted subunit A site, where the conformation of β-**maltoheptaose** bound via Glc7 O2ʹ, O1ʹ would result in unfavourable crystal contacts and where both Glc1 and Glc2 are closely similar to those in the **maltose**-bound structure (**19**) (Fig. **2B**). While the two alternative binding modes provide clear evidence of flexibility, the consistent recognition of alpha 1-4 linked glucose ligands (2-4 sugars) via Glc1 O3ʹ, O4ʹ suggests that the β-**maltoheptaose** O2ʹ, O1ʹ bound conformation in subunits B and C may arise from favourable crystal contacts, relative to the O3ʹ, O4ʹ bound conformation, rather than from a preference for βGlc7 O2ʹ, O1ʹ. In the β-**laminaritriose** structure the bound Glc3 orientation suggests that the remaining two glucose residues, which are not defined in the electron density, extend from the binding site above Glu321 and Asn323 and away from the surface of the protein in a previously unseen direction for small glucosyl-ligands (**7,19**). In each of the β-**isomaltotriose**, β-**laminaritriose** and β-**maltoheptaose** structures there are relatively long protein ligand interactions between Arg343 and the bound glucose O6ʹ while β-**isomaltotriose** exhibits multiple, significant interactions (2.96-3.32 Å) between Arg343 NH1/NH2 and O5 of the bound reducing terminal β-glucose (Table **3**) which are not present in any of the other structures but are mirrored in the galactose-bound structure (**7**). β-M**altoheptaose** and β-**laminaritriose** exhibit relatively weak binding (IC_50_ 3.43, 4.02 respectively) while β-**isomaltotriose** has high affinity binding (IC_50_ 1.99), comparable to that of **ManNAc** (IC_50_ 1.67).

The rfhSP-D bound structure of the dextran-based β-**isomaltotriose** ligand (Fig. **2F**) reveals for the first time an extended binding site for internal, non-terminal sugars beyond the calcium binding site where coordination is achieved via the O2ʹ and O1ʹ hydroxyls of the reducing terminal βGlc3. The ligand extends out of the calcium binding site (Glc3) in the direction of Arg343 with the central glucose Glc2 anchored to the protein surface via Glu347 and Arg349 of the protein surface. Sequence analysis reveals that these two side chains are conserved across many of the collectins, Glu347 in SP-D and Arg349 in both SP-D and the closely related surfactant protein A, which suggests that this binding surface may be important for longer ligand recognition across the collectin family. Indeed, structural studies of *S. enterica* Minnesota R5 and R7 oligosaccharides provide evidence for recognition of the inner core KdoIII, in addition to the calcium dependent binding of HepI, at this site (**9**). This site has also been implicated in the binding of α1-2 dimannose to hSP-D R343V (**6**). The electron density for the third glucose (Glc1) in the β-**isomaltotriose** chain suggests water-mediated interactions with a polar pocket beyond Glu347 and Arg349 and the anchored Glc2. The weak affinity of rfhSP-D for galactose (IC_50_ 12, Fig. **1**; Table **2**) is consistent with the secondary β-**isomaltotriose** site being the major determinant of high affinity β-**isomaltotriose** binding (IC_50_ 1.99), rather than the multiple, significant interactions of O5 with Arg343 NH1/NH2 which are also present for **galactose**. Of all the ligands studied here, β-**cellotriose,** bound via the non-reducing terminal Glc O3ʹ, O4ʹ in all three subunits although O2ʹ, O1ʹ is available at the reducing terminal, has the shortest distances to Arg343 (Fig. **2E****, 4B;** Table **3**) but exhibits only a moderate affinity (IC_50_ 2.81) for rfhSP-D (Fig. **1**; Table **2**). As reported by Crouch *et al*. (**19**), while direct bonding of Arg343 to bound ligand is stereochemically unlikely, the charged guanidyl group probably contributes to ligand stabilisation and may influence orientation.

While binding studies of β-**maltotetraose** have not been carried out here, previous work reports a moderately high affinity with an IC_50_ of 1.8 compared to maltose with an IC_50_ of 2.3 (both taken from Crouch *et al*. (**19**). At first sight this is consistent with the subunit A structure, where the non-Ca bound reducing terminal sugar returns to and interacts with the protein surface, as for β-**maltotriose** which demonstrates high affinity binding in all three subunits due to the secondary site involving Phe335 (**19**). However, the free conformation of β-**maltotetraose** in the sufficiently open binding site in subunits B and C, again bound via the non-reducing terminal O3ʹ, O4ʹ but extending away from the surface with only the bound terminal Glc visible in the electron density, indicates that this is the preferred conformation which will be reflected in binding studies. It seems likely that this preference for the free conformation of the β-**maltotetraose** is due to a weaker binding affinity for the somewhat strained conformation despite the secondary interaction. Were Phe335 to indeed provide a physiologically meaningful secondary binding site for the third sugar in extended maltopolysaccharides (**19**), rather than reflecting only the ability of the non-Ca bound reducing terminal sugar of β-**maltotriose** to return to and interact with the protein surface, then the same interaction would be expected to be preferred for both β-**maltotetraose** and β-**maltoheptaose**, which is not the case.

β**-laminaritriose and** β**-cellotriose**, the first poly β-D-glucose ligands to be investigated bound to rfhSP-D, are important as analogues for β-glucan-based and cellulose-based ligands such as the cell wall of fungi and allergens that have been shown to be recognised by SP-D (**25,26**). Across all three subunits these two β-linked ligands bind via opposite ends of the trisaccharide and present the opposite face of the bound Glc to Asp325, with β**-laminaritriose** (β1-3 linked) bound via βGlc3 O2ʹ, O1ʹ (reducing) and β**-cellotriose** (β1-4 linked) via βGlc1 O3ʹ, O4ʹ, suggesting that the differing β-linkages influence the preferred binding mode.

The mode of carbohydrate recognition by hSP-D is difficult to predict (**7,20,23,24**), depending as it does on a variety of parameters including the carbohydrate itself, the nature of the glycosidic linkage(s), interaction with the binding site flanking residues Arg343 and Asp325 (**6,7,9,17,27**), and the possibility of utilising a supplementary binding site for sugars which extend from and beyond the calcium binding site as shown here for β-**isomaltotriose** and by (**9**). It is clear however that provided a pair of hydroxyls in a mannose-type O3ʹ, O4ʹ configuration is available then hSP-D has the versatility not only to recognise a wide variety of carbohydrate ligands, but to adopt alternative mechanisms to recognise the same ligand, as demonstrated for bacterially derived LPS (**9**), synthetic oligosaccharides (**17**) and β-**maltoheptaose** as reported here.

The interaction of SP-D with β-D-glucose, via the O2ʹ and O1ʹ hydroxyls at the saccharide reducing terminal, may be important in recognising ligands released from the surface of pathogens into the extracellular milieu (**28**), or, in the case of bacterial LPS, recognition of the reducing terminal of the O antigen prior to attachment to the separately synthesised core-lipid A and subsequent presentation of the mature LPS at the bacterial outer membrane surface (**29**). β-**cellotriose**, β-**isomaltotriose** and β-**laminaritriose** are significant as analogues for cellulose, dextran and β-glucan-based ligands respectively, not only as the exo-polysaccharides of bacteria, fungi and allergens (**30**) but also in their own right. This may be of some significance in dextran-mediated drug delivery (**31**) and the role of β-glucans as immune stimulants (**32**).

## EXPERIMENTAL PROCEDURES

### The recombinant fragment of human SP-D (rfhSP-D)

A recombinant homotrimeric fragment of human SP-D (rfhSP-D) was expressed in bacteria and purified as described previously (**33**), or from the culture supernatant of a stable transfected Chinese Hamster Ovary cell line and purified using affinity chromatography with an N-acetylmannosamine (ManNAc)-coupled matrix as described previously (**34**). Each chain contained 177 amino acids (Gly179 to Phe355) comprised of a short collagenous region with eight Gly-Xaa-Yaa repeats, the α-helical coiled coil neck region, and the globular CRD. The endotoxin level was measured by the *Limulus* Amebocyte Lysate Assay (ThermoFisher Scientific) and preparations contained less than 5 pg endotoxin/µg rfhSP-D.

### Competitive mannan-based enzyme linked immunosorbent assay (ELISA)

To assess the binding of the recombinant fragment of human SP-D (rfhSP-D) to different carbohydrates, a competitive mannan-based enzyme linked immunosorbent assay was implemented. For this, Maxisorp 96 well microtitre plates (Nunc) were coated with 100 µL of mannan (50 µg/mL) (Sigma) in 0.1M sodium carbonate-bicarbonate coating buffer (15 mM Na_2_CO_3_, 35 mM NaHCO_3_, pH 9.6) and incubated overnight at 4°C. The plates were then washed 4 times using wash buffer (50 mM Tris-HCL, 150 mM NaCl pH 7.5 TBS, 5 mM calcium chloride, 0.05% v/v Tween 20) before blocking using a 2% w/v bovine serum albumin (BSA) in wash buffer solution for 1 hour at room temperature. Different carbohydrates solutions were prepared in wash buffer and diluted two-fold, each containing 10 µg/mL of rfhSP-D for experiments used to test calcium-dependent binding. For samples controlling for non-calcium dependent binding, 10 mM EDTA (Sigma) was added to the diluent mix. The sugars were tested in the following concentration range: β-Galactose dilution range: 250-0.25 mM; Maltose, ManNAc, and β-Maltoheptaose dilution range: 100-0.1 mM; β-Maltotriose, β-Isomaltotriose, β-Laminaritriose, β-Cellotriose dilution range: 50-0.05 mM. All carbohydrates were purchased from Sigma except β-Cellotriose, β-Maltotriose, and β-Laminaritriose which were purchased from Megazyme. After 1 hour incubation with continuous stirring, the carbohydrate-rfhSP-D solutions were added to the blocked plates (100 µL per well) and incubated overnight at 4°C. The bound rfhSP-D was detected by sequential 1 hour incubations with a rabbit polyclonal anti rfhSP-D antibody (stock concentration 1.61 mg/mL, final dilution 1:1000 in wash buffer) followed by horseradish peroxidase-conjugated goat anti-rabbit IgG antibody (Invitrogen, 1:2000 in wash buffer) at room temperature. At the end of the incubation, the plates were washed as previously described and developed using 100 µL/well of 5,5’-tetremathylbenzidine (TMB) substrate (Thermo Scientific). The colorimetric reaction was stopped by adding 50 µL of 0.5 M H_2_SO_4_ to each well. The absorbance was measured using a FLUOstar Omega plate reader (BMG Labtech) at 450 nm. A sigmoidal 4 parameter logistic curve was fitted to each absorbance curve and the half-maximal inhibitory concentration showing the concentration of carbohydrate required to inhibit 50% of rfhSP-D binding to mannan (IC_50_) was calculated using GraphPad Prism (version 10). This value was used to determine binding affinity of the different carbohydrates to rfhSP-D.

### Crystallisation and data collection

The recombinant homotrimeric fragment of human SP-D (rfhSP-D) at 1 mg/ml was dialysed into 10 mM Tris, 10 mM calcium chloride, 140 mM NaCl, 0.02% NaN_3_ at pH7.5 and concentrated to 7.7 mg/ml.

Native crystals of rfhSP-D were grown in sitting drops using screens around conditions previously reported (**21**). For β-maltoheptaose (M7) and β-maltotetraose (M4) ligand was soaked into native crystals grown in 0.1 M Tris pH 7.5, 16% PEG 6000 and 0.1 M Tris, pH 8.0, 16% PEG 4000 respectively. The ligand was added to cryoprotectant to a final concentration of 10 mM in the drop. The crystals were prepared for cryocooling by successive addition of increasing concentration MPD (5-20%) in 2 µl aliquots over around half an hour and exchanging 8 µl of the 20% solution. 10 mM calcium chloride was added to the cryoprotectant.

β-isomaltotriose (I3) was soaked into native crystals grown in 0.1 M Tris pH 6.0, 16% PEG 6000, 10 mM calcium chloride. Additional calcium was included in the cryoprotectant, which included 20 mM ligand, to maintain the calcium concentration during cryoprotection. β-cellotriose (C3) was soaked into native rfhSP-D crystals grown in 0.1 M Tris pH 6.0, 16% PEG 6000, with 10 mM calcium chloride and 20 mM ligand added to the cryoprotectant in a similar way to β-isomaltotriose.

β-laminaritriose (L3) was initially soaked into native crystals grown in 0.1 M Tris pH 6.0, 16% PEG 6000, 10 mM calcium chloride during cryoprotection with MPD and 20mM ligand, however the crystals dissolved during protection. The crystals later reformed resulting in a co-crystallised structure with rfhSP-D. Additional cryoprotection buffer (MPD) was added to ensure suitability for cryocooling.

β-maltoheptaose data were collected on station 14.1 and β-maltotetraose data were collected on station 10.1 at Daresbury SRS. β-isomaltotriose, β-cellotriose and β-laminaritriose data were collected on i04-1, i02 and i04-1, respectively, at Diamond Light Source. Integrated intensities were calculated using *iMOSFLM* (**35**) and data was processed using the *Aimless*, *Truncate*, *Uniquify*, and *Sortmtz* programmes as part of the CCP4 programme suite (**36**). Data collection and processing statistics are given in Table **5**.

**Table 5.**
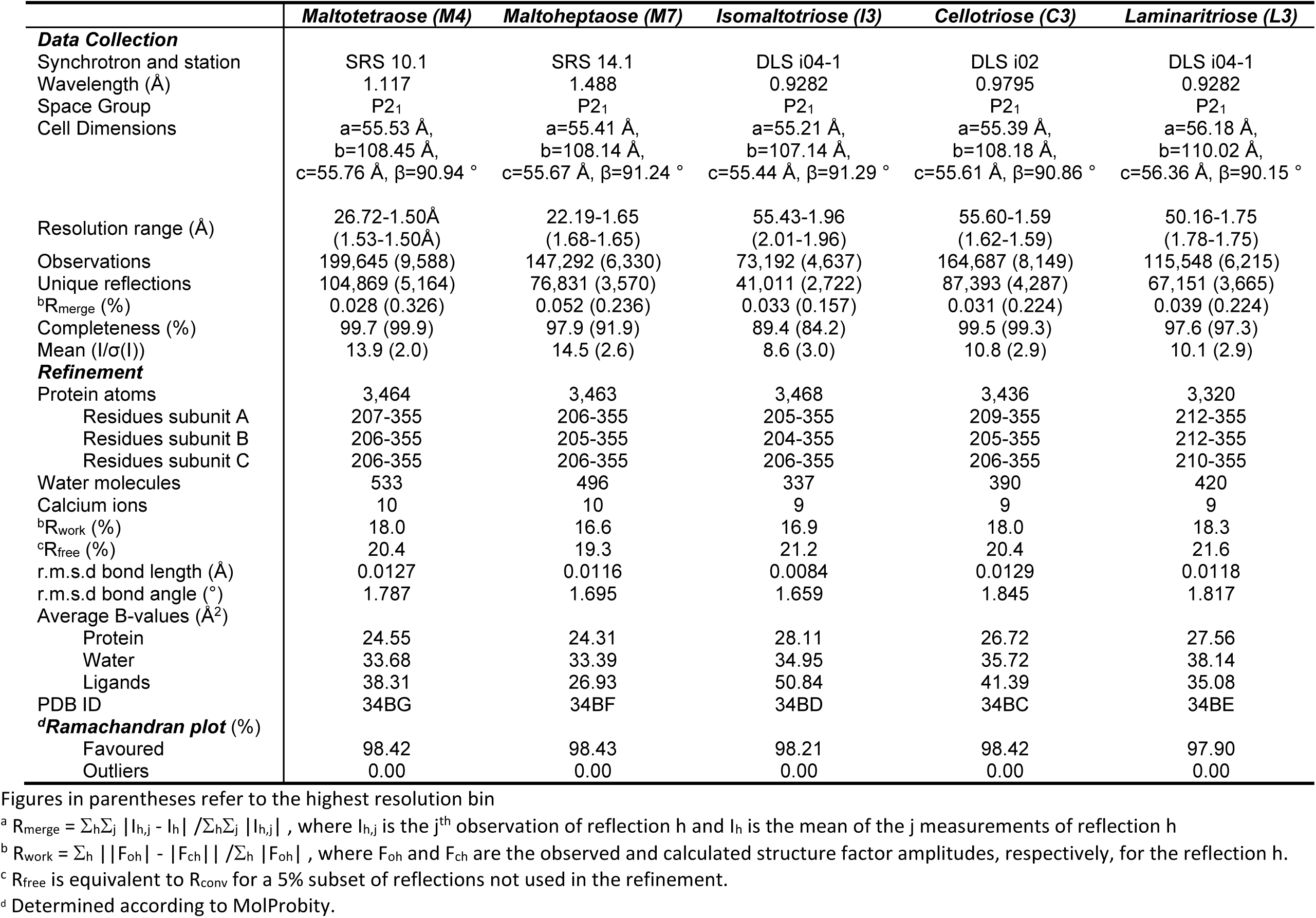
Data collection and processing.

|  | ***Maltotetraose (M4)*** | ***Maltoheptaose (M7)*** | ***Isomaltotriose (I3)*** | ***Cellotriose (C3)*** | ***Laminaritriose (L3)*** |
|---|---|---|---|---|---|
| ***Data Collection*** |  |  |  |  |  |
| Synchrotron and station | SRS 10.1 | SRS 14.1 | DLS i04-1 | DLS i02 | DLS i04-1 |
| Wavelength (Å) | 1.117 | 1.488 | 0.9282 | 0.9795 | 0.9282 |
| Space Group | P2<sub>1</sub> | P2<sub>1</sub> | P2<sub>1</sub> | P2<sub>1</sub> | P2<sub>1</sub> |
| Cell Dimensions | a=55.53 Å,<br>b=108.45 Å,<br>c=55.76 Å, β=90.94 ° | a=55.41 Å,<br>b=108.14 Å,<br>c=55.67 Å, β=91.24 ° | a=55.21 Å,<br>b=107.14 Å,<br>c=55.44 Å, β=91.29 ° | a=55.39 Å,<br>b=108.18 Å,<br>c=55.61 Å, β=90.86 ° | a=56.18 Å,<br>b=110.02 Å,<br>c=56.36 Å, β=90.15 ° |
| Resolution range (Å) | 26.72-1.50Å<br>(1.53-1.50Å) | 22.19-1.65<br>(1.68-1.65) | 55.43-1.96<br>(2.01-1.96) | 55.60-1.59<br>(1.62-1.59) | 50.16-1.75<br>(1.78-1.75) |
| Observations | 199,645 (9,588) | 147,292 (6,330) | 73,192 (4,637) | 164,687 (8,149) | 115,548 (6,215) |
| Unique reflections | 104,869 (5,164) | 76,831 (3,570) | 41,011 (2,722) | 87,393 (4,287) | 67,151 (3,665) |
| <sup>b</sup>R<sub>merge</sub> (%) | 0.028 (0.326) | 0.052 (0.236) | 0.033 (0.157) | 0.031 (0.224) | 0.039 (0.224) |
| Completeness (%) | 99.7 (99.9) | 97.9 (91.9) | 89.4 (84.2) | 99.5 (99.3) | 97.6 (97.3) |
| Mean (I/σ(I)) | 13.9 (2.0) | 14.5 (2.6) | 8.6 (3.0) | 10.8 (2.9) | 10.1 (2.9) |
| ***Refinement*** |  |  |  |  |  |
| Protein atoms | 3,464 | 3,463 | 3,468 | 3,436 | 3,320 |
| Residues subunit A | 207-355 | 206-355 | 205-355 | 209-355 | 212-355 |
| Residues subunit B | 206-355 | 205-355 | 204-355 | 205-355 | 212-355 |
| Residues subunit C | 206-355 | 206-355 | 206-355 | 206-355 | 210-355 |
| Water molecules | 533 | 496 | 337 | 390 | 420 |
| Calcium ions | 10 | 10 | 9 | 9 | 9 |
| <sup>b</sup>R<sub>work</sub> (%) | 18.0 | 16.6 | 16.9 | 18.0 | 18.3 |
| <sup>c</sup>R<sub>free</sub> (%) | 20.4 | 19.3 | 21.2 | 20.4 | 21.6 |
| r.m.s.d bond length (Å) | 0.0127 | 0.0116 | 0.0084 | 0.0129 | 0.0118 |
| r.m.s.d bond angle (°) | 1.787 | 1.695 | 1.659 | 1.845 | 1.817 |
| Average B-values (Å<sup>2</sup>) |  |  |  |  |  |
| Protein | 24.55 | 24.31 | 28.11 | 26.72 | 27.56 |
| Water | 33.68 | 33.39 | 34.95 | 35.72 | 38.14 |
| Ligands | 38.31 | 26.93 | 50.84 | 41.39 | 35.08 |
| PDB ID | 34BG | 34BF | 34BD | 34BC | 34BE |
| ***<sup>d</sup>Ramachandran plot*** (%) |  |  |  |  |  |
| Favoured | 98.42 | 98.43 | 98.21 | 98.42 | 97.90 |
| Outliers | 0.00 | 0.00 | 0.00 | 0.00 | 0.00 |

Figures in parentheses refer to the highest resolution bin
<sup>a</sup> R<sub>merge</sub> = Σ<sub>h</sub>Σ<sub>j</sub> |I<sub>h,j</sub> - I<sub>h</sub>| /Σ<sub>h</sub>Σ<sub>j</sub> |I<sub>h,j</sub>| , where I<sub>h,j</sub> is the j<sup>th</sup> observation of reflection h and I<sub>h</sub> is the mean of the j measurements of reflection h
<sup>b</sup> R<sub>work</sub> = Σ<sub>h</sub> ||F<sub>oh</sub>| - |F<sub>ch</sub>|| /Σ<sub>h</sub> |F<sub>oh</sub>| , where F<sub>oh</sub> and F<sub>ch</sub> are the observed and calculated structure factor amplitudes, respectively, for the reflection h.
<sup>c</sup> R<sub>free</sub> is equivalent to R<sub>conv</sub> for a 5% subset of reflections not used in the refinement.
<sup>d</sup> Determined according to MolProbity.

### Crystallographic structure solution and refinement

All structures were solved by molecular replacement using the previously determined 1.6 Å native rfhSP-D structure (**21**) for the ligand-bound rfhSP-D structures. The structures were refined using the program *Refmac5* (**37**). The *CCP4 program suite* (36,38) was used to calculate maps and models built using the graphics program *COOT* (**39**). Topology and parameter files for ligands were already present in the CCP4 reference dictionaries. Refinement statistics are shown in Table **5**. Structure quality was checked using *MolProbity* (**40**) and *Privateer* as part of the CCP4i2 suite (**41**) as well as the Protein Data Bank (PDB) validation software. The expression cassette contains a short stretch of 8 Gly-X-Y repeats leading into the neck, which starts at residue 203, which are not seen in the electron density. The N-terminal neck region itself is flexible and seen to varying degrees in the electron density, as is the case for other SP-D structures. For each subunit in each structure the cutoff (see Table **5**) as to where amino acids have been fitted into the density has been made based on both the density present, particularly for main chain, and the validation statistics. All main and side-chain stereochemical parameters are within accepted limits or better, with more than 97% of residues in favoured regions of the Ramachandran plot with no outliers. Final refinement statistics are provided in Table **5**. Molecular figures were generated using CCP4mg molecular graphics software (**42**).

## Data availability

The coordinates and structure factors for the cellotriose (34BC), isomaltotriose (34BD), laminaritriose (34BE), maltoheptaose (34BF), and maltotetraose (34BG) ligand-bound structures have been deposited with the PDB. The raw experimental diffraction image data are available *via* Keele University Data Repository, DOI: 10.21252/2d2f-v394, 10.21252/2naf-vz07, 10.21252/tt63-9e76, 10.21252/21j8-2f16, 10.21252/jjc5-qh80 for 34BC, 34BD, 34BE, 34BF, and 34BG respectively.

### Supporting information

This article contains supporting information.

## Acknowledgements

We thank the beamline scientists at the Daresbury SRS and Diamond Light Source for their help and support.

## Author contributions

J.R.L., T.C-H. and A.J.S. Writing – original draft; J.R.L., T.C-H., H.M.W., A.J.S., H.W.C., A.W., J.M., T.J.G. and A.K.S. Writing – review & editing; J.R.L., T.C-H., H.M.W., A.J.S., Y.G., G.B., A.W., I.R.B., T.J.G. and A.K.S. Methodology; J.R.L., T.C-H., H.M.W., A.J.S., Y.G., G.B., A.W., I.R.B., T.J.G. and A.K.S. Investigation; J.R.L., T.C-H., H.M.W., A.J.S., Y.G., G.B., A.W., I.R.B., T.J.G. and A.K.S. Formal analysis; H.M.W., T.C-H. and A.K.S. Data curation; H.M.W., J.M., T.C-H., A.W., A.K.S. and T.J.G. Visualisation; H.M.W., J.M., T.C-H., A.W., A.K.S. and T.J.G. Validation; A.W., J.M., H.W.C., T.C-H., G.B., K.B.M.R., A.K.S. and T.J.G. Funding acquisition; J.M., H.W.C., T.J.G., K.B.M.R. and A.K.S. Resources; J.M., H.W.C., T.J.G., K.B.M.R., A.W. and A.K.S. Project administration; K.B.M.R., T.J.G., A.K.S., H.W.C. and J.M. Conceptualisation; A.K.S. Supervision

## Funding and additional information

This work was supported by Science and Technology Facilities Council, and Diamond Light Source Awards MX14692 and MX8359 (T.J.G, A.K.S), and by MRC award G0400743 (A.K.S, T.J.G). J.M, T.C.H. and H.W.C would like to thank the Medical Research Council Catalyst for the Developmental Pathway Funding Scheme (DPFS) Award MR/P026907/1. G.B. thanks the UCL MRC Doctoral Training Programme for funding her PhD and A.W. the University of Southampton MRC Doctoral Training Programme for funding his PhD. J.R.L., H.M.W. and A.J.S. thank the University of Keele for PhD funding support.

## Declaration of competing interest

The authors declare that they have no known competing financial interests or personal relationships that could have appeared to influence the work reported in this paper.

## Data availability

### Author contributions

J.R.L., T.C-H. and A.J.S. Writing-original draft; J.R.L., T.C-H., H.M.W., A.J.S., H.W.C., A.W., J.M., T.J.G. and A.K.S. Writing-review & editing; J.R.L., T.C-H., H.M.W., A.J.S., Y.G., G.B., A.W., I.R.B., T.J.G. and A.K.S. Methodology; J.R.L., T.C-H., H.M.W., A.J.S., Y.G., G.B., A.W., I.R.B., T.J.G. and A.K.S. Investigation; J.R.L., T.C-H., H.M.W., A.J.S., Y.G., G.B., A.W., I.R.B., T.J.G. and A.K.S. Formal analysis; H.M.W., T.C-H. and A.K.S. Data curation; H.M.W., J.M., T.C-H., A.W., A.K.S. and T.J.G. Visualisation; H.M.W., J.M., T.C-H., A.W., A.K.S. and T.J.G. Validation; A.W., J.M., H.W.C., T.C-H., G.B., K.B.M.R., A.K.S. and T.J.G. Funding acquisition; J.M., H.W.C., T.J.G., K.B.M.R. and A.K.S. Resources; J.M., H.W.C., T.J.G., K.B.M.R., A.W. and A.K.S. Project administration; K.B.M.R., T.J.G., A.K.S., H.W.C. and J.M. Conceptualisation; A.K.S. Supervision

## Notes

https://researchdata.keele.ac.uk/140/

https://researchdata.keele.ac.uk/142/

https://researchdata.keele.ac.uk/144/

https://researchdata.keele.ac.uk/145/

### Competing Interest Statement

The authors have declared no competing interest.

## References

1. Ujma, S., Horsnell, W. G., Katz, A. A., Clark, H. W., and Schäfer, G. (2017) Non-Pulmonary Immune Functions of Surfactant Proteins A and D. J. Innate Immun. 9, 3–11. 10.1159/000451026

2. Reid K. B. (1998) Interactions of surfactant protein D with pathogens, allergens and phagocytes. Biochim. Biophys. Acta. 1408, 290–295. 10.1016/s0925-4439(98)00074-x

3. LeVine, A. M., and Whitsett, J. A. (2001) Pulmonary collectins and innate host defense of the lung. Microbes Infect. 3, 161–166. 10.1016/s1286-4579(00)01363-0

4. Clark, H., and Reid, K. B. (2002) Structural requirements for SP-D function in vitro and in vivo: therapeutic potential of recombinant SP-D. Immunobiology 205, 619–631. 10.1078/0171-2985-00159

5. Singh, M., Madan, T., Waters, P., Parida, S. K., Sarma, P. U., and Kishore, U. (2003) Protective effects of a recombinant fragment of human surfactant protein D in a murine model of pulmonary hypersensitivity induced by dust mite allergens. Immunol Lett. 86, 299–307. 10.1016/s0165-2478(03)00033-6

6. Crouch, E., Hartshorn, K., Horlacher, T., McDonald, B., Smith, K., Cafarella, T., Seaton, B., Seeberger, P. H., and Head, J. (2009) Recognition of mannosylated ligands and influenza A virus by human surfactant protein D: contributions of an extended site and residue 343. Biochemistry 48, 3335–3345. 10.1021/bi8022703

7. Shrive, A. K., Martin, C., Burns, I., Paterson, J. M., Martin, J. D., Townsend, J. P., Waters, P., Clark, H. W., Kishore, U., Reid, K. B., and Greenhough, T. J. (2009) Structural characterisation of ligand-binding determinants in human lung surfactant protein D: influence of Asp325. J. Mol. Biol. 394, 776–788. 10.1016/j.jmb.2009.09.057

8. Clark, H. W., Mackay, R. M., Deadman, M. E., Hood, D. W., Madsen, J., Moxon, E. R., Townsend, J. P., Reid, K. B. M., Ahmed, A., Shaw, A. J., Greenhough, T. J., and Shrive, A. K. (2016) Crystal Structure of a Complex of Surfactant Protein D (SP-D) and Haemophilus influenzae Lipopolysaccharide Reveals Shielding of Core Structures in SP-D-Resistant Strains. Infect. Immun. 84, 1585–1592. 10.1128/IAI.01239-15

9. Littlejohn, J. R., da Silva, R. F., Neale, W. A., Smallcombe, C. C., Clark, H. W., Mackay, R. A., Watson, A. S., Madsen, J., Hood, D. W., Burns, I., Greenhough, T. J., and Shrive, A. K. (2018) Structural definition of hSP-D recognition of Salmonella enterica LPS inner core oligosaccharides reveals alternative binding modes for the same LPS. PLoS one 13, e0199175. 10.1371/journal.pone.0199175

10. Serra, D. O., Richter, A. M., and Hengge, R. (2013) Cellulose as an architectural element in spatially structured *Escherichia coli* biofilms. J. bacteriol. 195, 5540–5554. 10.1128/JB.00946-13

11. Vu, B., Chen, M., Crawford, R. J., and Ivanova, E. P. (2009) Bacterial extracellular polysaccharides involved in biofilm formation. Molecules 14, 2535–2554. 10.3390/molecules14072535

12. McIntosh, M., Stone, B. A., and Stanisich, V. A. (2005) Curdlan and other bacterial (1-->3)-beta-D-glucans. Appl. Microbiol. Biotechnol. 68, 163–173. 10.1007/s00253-005-1959-5

13. Fakih, D., Pilecki, B., Schlosser, A., Jepsen, C. S., Thomsen, L. K., Ormhøj, M., Watson, A., Madsen, J., Clark, H. W., Barfod, K. K., Hansen, S., Marcussen, N., Jounblat, R., Chamat, S., Holmskov, U., and Sorensen, G. L. (2015) Protective effects of surfactant protein D treatment in 1,3-β-glucan-modulated allergic inflammation. Am. J. Physiol. Lung Cell Mol. Physiol. 309, L1333–L1343. 10.1152/ajplung.00090.2015

14. Taylor, L. P., and Hepler, P. K. (1997) POLLEN GERMINATION AND TUBE GROWTH. Annu. Rev. Plant Physiol. Plant Mol. Biol. 48, 461–491. 10.1146/annurev.arplant.48.1.461

15. Nakanishi, I., Kimura, K., Suzuki, T., Ishikawa, M., Banno, I., Sakane, T., and Harada, T. (1976) Demonstration of curdlan-type polysaccharide and some other β-1,3-glucan in microorganisms with aniline blue. J. General Applied Microbiol. 22, 1–11. 10.2323/jgam.22.1

16. Sutherland, I.W. (2001) Microbial polysaccharides from Gram-negative bacteria. International Dairy Journal 11, 663–674. 10.1016/S0958-6946(01)00112-1

17. Williams, H. M., Watson, A., Madsen, J., Clark, H. W., Hood, D. W., Oscarson, S., Greenhough, T. J., and Shrive, A. K. (2026) Phosphorylation of inner core heptose is a major determinant of bacterial surface lipopolysaccharide recognition by the innate immune protein hSP-D. J. Biol. Chem. 302, 111307. 10.1016/j.jbc.2026.111307

18. Goh, B. C., Rynkiewicz, M. J., Cafarella, T. R., White, M. R., Hartshorn, K. L., Allen, K., Crouch, E. C., Calin, O., Seeberger, P. H., Schulten, K., and Seaton, B. A. (2013) Molecular mechanisms of inhibition of influenza by surfactant protein D revealed by large-scale molecular dynamics simulation. Biochemistry 52, 8527–8538. 10.1021/bi4010683

19. Crouch, E., McDonald, B., Smith, K., Cafarella, T., Seaton, B., and Head, J. (2006a) Contributions of phenylalanine 335 to ligand recognition by human surfactant protein D: ring interactions with SP-D ligands. J. Biol. Chem. 281, 18008–18014. 10.1074/jbc.M601749200

20. Allen, M. J., Laederach, A., Reilly, P. J., and Mason, R. J. (2001) Polysaccharide recognition by surfactant protein D: novel interactions of a C-type lectin with nonterminal glucosyl residues. Biochemistry 40, 7789–7798. 10.1021/bi002901q

21. Shrive, A. K., Tharia, H. A., Strong, P., Kishore, U., Burns, I., Rizkallah, P. J., Reid, K. B., and Greenhough, T. J. (2003) High-resolution structural insights into ligand binding and immune cell recognition by human lung surfactant protein D. J. Mol. Biol. 331, 509–523. 10.1016/s0022-2836(03)00761-7

22. Crouch, E. C., Smith, K., McDonald, B., Briner, D., Linders, B., McDonald, J., Holmskov, U., Head, J., and Hartshorn, K. (2006b) Species differences in the carbohydrate binding preferences of surfactant protein D. Am. J. Respir. Cell. Mol. Biol. 35, 84–94. 10.1165/rcmb.2005-0462OC

23. Ng, K. K., Drickamer, K., and Weis, W. I. (1996) Structural analysis of monosaccharide recognition by rat liver mannose-binding protein. J. Biol. Chem. 271, 663–674. 10.1074/jbc.271.2.663

24. Ng, K. K., Kolatkar, A. R., Park-Snyder, S., Feinberg, H., Clark, D. A., Drickamer, K., and Weis, W. I. (2002) Orientation of bound ligands in mannose-binding proteins. Implications for multivalent ligand recognition. J. Biol. Chem. 277, 16088–16095. 10.1074/jbc.M200493200

25. Madan, T., Kishore, U., Singh, M., Strong, P., Clark, H., Hussain, E. M., Reid, K. B., and Sarma, P. U. (2001) Surfactant proteins A and D protect mice against pulmonary hypersensitivity induced by Aspergillus fumigatus antigens and allergens. J. Clin. Invest. 107, 467–475. 10.1172/JCI10124

26. Erpenbeck, V. J., Malherbe, D. C., Sommer, S., Schmiedl, A., Steinhilber, W., Ghio, A. J., Krug, N., Wright, J. R., and Hohlfeld, J. M. (2005) Surfactant protein D increases phagocytosis and aggregation of pollen-allergen starch granules. Am. J. Physiol. Lung Cell Mol. Physiol. 288, L692–L698. 10.1152/ajplung.00362.2004

27. Crouch, E., McDonald, B., Smith, K., Roberts, M., Mealy, T., Seaton, B., and Head, J. (2007) Critical role of Arg/Lys343 in the species-dependent recognition of phosphatidylinositol by pulmonary surfactant protein D. Biochemistry 46, 5160–5169. 10.1021/bi700037x

28. Digby, J., Kalbfleisch, J., Glenn, A., Larsen, A., Browder, W., and Williams, D. (2003) Serum glucan levels are not specific for presence of fungal infections in intensive care unit patients. Clin. Diagn. Lab. Immunol. 10, 882–885. 10.1128/cdli.10.5.882-885.2003

29. Raetz, C. R., Reynolds, C. M., Trent, M. S., and Bishop, R. E. (2007) Lipid A modification systems in gram-negative bacteria. Annu. Rev. Biochem. 76, 295–329. 10.1146/annurev.biochem.76.010307.145803

30. Crouch E. C. (1998) Collectins and pulmonary host defense. Am. J. Respir. Cell Mol. Biol. 19, 177–201. 10.1165/ajrcmb.19.2.140

31. Varshosaz J. (2012) Dextran conjugates in drug delivery. Expert Opin. Drug Deliv. 9, 509–523. 10.1517/17425247.2012.673580

32. Akramiene, D., Kondrotas, A., Didziapetriene, J., and Kevelaitis, E. (2007) Effects of beta-glucans on the immune system. *Medicina (Kaunas*, Lithuania), 43, 597–606. https://pubmed.ncbi.nlm.nih.gov/17895634/

33. Strong, P., Reid, K. B., and Clark, H. (2002) Intranasal delivery of a truncated recombinant human SP-D is effective at down-regulating allergic hypersensitivity in mice sensitized to allergens of Aspergillus fumigatus. Clin. Exp. Immunol. 130, 19–24. 10.1046/j.1365-2249.2002.01968.x

34. Sorensen, G. L., Hoegh, S. V., Leth-Larsen, R., Thomsen, T. H., Floridon, C., Smith, K., Kejling, K., Tornoe, I., Crouch, E. C., and Holmskov, U. (2009) Multimeric and trimeric subunit SP-D are interconvertible structures with distinct ligand interaction. Mol. Immunol. 46, 3060–3069. 10.1016/j.molimm.2009.06.005

35. Battye, T. G., Kontogiannis, L., Johnson, O., Powell, H. R., and Leslie, A. G. (2011) iMOSFLM: a new graphical interface for diffraction-image processing with MOSFLM. Acta Crystallogr. D Biol. Crystallogr. 67, 271–281. 10.1107/S0907444910048675

36. Winn, M. D., Ballard, C. C., Cowtan, K. D., Dodson, E. J., Emsley, P., Evans, P. R., Keegan, R. M., Krissinel, E. B., Leslie, A. G., McCoy, A., McNicholas, S. J., Murshudov, G. N., Pannu, N. S., Potterton, E. A., Powell, H. R., Read, R. J., Vagin, A., and Wilson, K. S. (2011) Overview of the CCP4 suite and current developments. Acta Crystallogr. D Biol. Crystallogr. 67, 235–242. 10.1107/S0907444910045749

37. Murshudov, G. N., Skubák, P., Lebedev, A. A., Pannu, N. S., Steiner, R. A., Nicholls, R. A., Winn, M. D., Long, F., and Vagin, A. A. (2011) REFMAC5 for the refinement of macromolecular crystal structures. Acta Crystallogr. D Biol. Crystallogr. 67, 355–367. 10.1107/S0907444911001314

38. Agirre, J., et al. (2023) The CCP4 suite: integrative software for macromolecular crystallography. Acta Crystallogr. D Biol. Crystallogr. 79, 449–461. 10.1107/S2059798323003595

39. Emsley, P., Lohkamp, B., Scott, W. G., and Cowtan, K. (2010) Features and development of Coot. Acta Crystallogr. D Biol. Crystallogr. 66, 486–501. 10.1107/S0907444910007493

40. Potterton, L., Agirre, J., Ballard, C., Cowtan, K., Dodson, E., Evans, P. R., Jenkins, H. T., Keegan, R., Krissinel, E., Stevenson, K., Lebedev, A., McNicholas, S. J., Nicholls, R. A., Noble, M., Pannu, N. S., Roth, C., Sheldrick, G., Skubak, P., Turkenburg, J., Uski, V., von Delft F., Waterman, D., Wilson, K., Winn, M., Wojdyr, M. (2018) CCP4i2: the new graphical user interface to the CCP4 program suite. Acta Crystallogr. D Biol. Crystallogr. 74, 68–84. 10.1107/S2059798317016035

41. Chen, V. B., Arendall, W. B., 3rd, Headd, J. J., Keedy, D. A., Immormino, R. M., Kapral, G. J., Murray, L. W., Richardson, J. S., and Richardson, D. C. (2010) MolProbity: all-atom structure validation for macromolecular crystallography. Acta Crystallogr. D Biol. Crystallogr. 66, 12–21. 10.1107/S0907444909042073

42. McNicholas, S., Potterton, E., Wilson, K. S., and Noble, M. E. (2011) Presenting your structures: the CCP4mg molecular-graphics software. Acta Crystallogr. D Biol. Crystallogr. 67, 386–394. 10.1107/S0907444911007281

