## Supplementary for "Recognition of β-D-glucose by surfactant protein D and structural evidence for an extended carbohydrate binding site"

**Figure S1. Competitive binding of rfhSP-D to mannan by different carbohydrates.**

**Figure S2. Competitive binding of rfhSP-D to different carbohydrates depends on the presence of Calcium ($Ca^{2+}$) in solution.**

**Table S1. Calcium-protein binding distances (Å).**

**Figure S1. Figure S1. Competitive binding of rfhSP-D to mannan by different carbohydrates.**
These values were obtained using a competitive mannan-based ELISA described in the methods section. Data shown as mean ± standard deviation of 2 technical repeats.

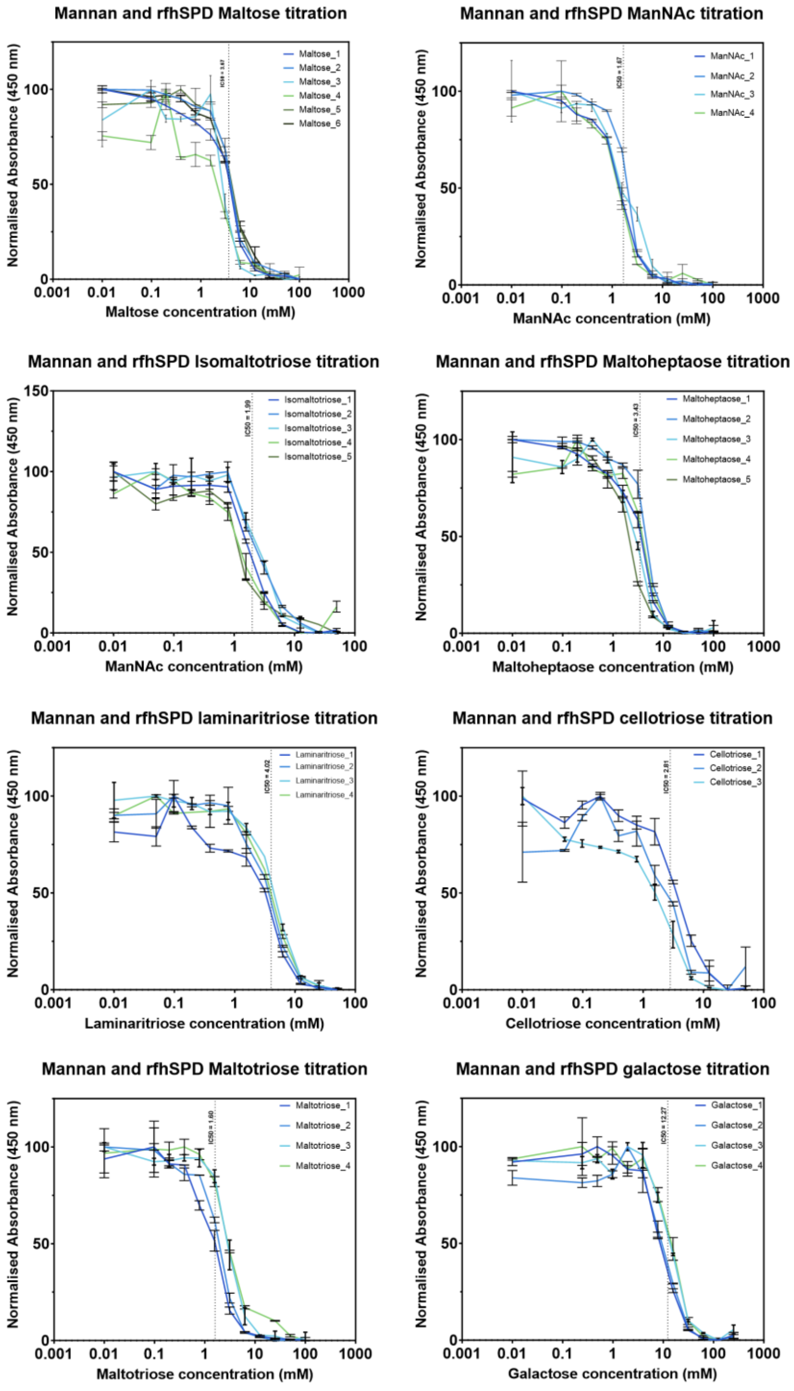

**Figure S2. Competitive binding of rfhSP-D to different carbohydrates depends on the presence of Calcium ($Ca^{2+}$) in solution.**
The values reported were obtained using competitive mannan ELISAs (described in methods section) in the presence of 5 mM calcium chloride (filled markers) and 10 mM EDTA (unfilled markers). Data shown as mean ± standard deviation of 2 technical repeats.

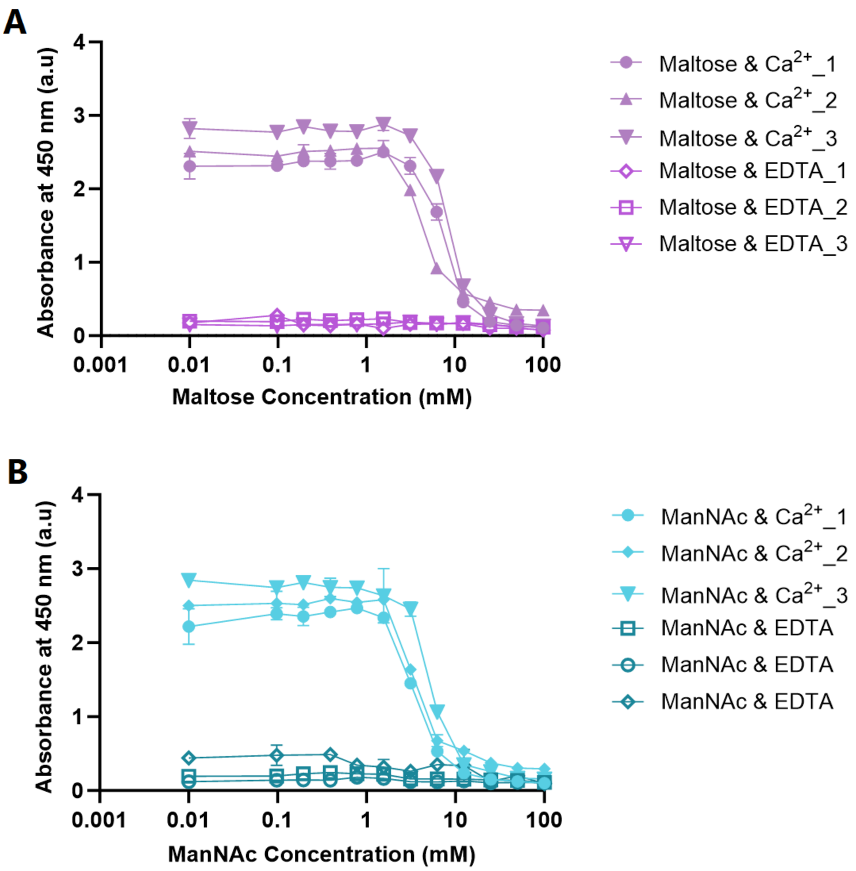

**Table S1. Calcium-protein binding distances (Å).**

| Atom 1 | Atom 2 | | | Maltotetraose (M4) | | | Maltoheptaose (M7) | | | Isomaltotriose (I3) | | | Cellotriose (C3) | | | Laminaritriose (L3) | | |
|---|---|---|---|---|---|---|---|---|---|---|---|---|---|---|---|---|---|---|
| | | | | A | B | C | A | B | C | A | B | C | A | B | C | A | B | C |
| Ca1 | Glu | 321 | OE1 | 2.55 | 2.51 | 2.52 | 2.50 | 2.53 | 2.49 | 2.53 | 2.70 | 2.62 | 2.61 | 2.56 | 2.59 | 2.64 | 2.60 | 2.61 |
| | Asn | 323 | OD1 | 2.40 | 2.39 | 2.37 | 2.40 | 2.38 | 2.39 | 2.45 | 2.48 | 2.53 | 2.39 | 2.36 | 2.33 | 2.46 | 2.46 | 2.44 |
| | Glu | 329 | OE1 | 2.48 | 2.42 | 2.42 | 2.45 | 2.43 | 2.39 | 2.40 | 2.36 | 2.32 | 2.41 | 2.43 | 2.38 | 2.44 | 2.40 | 2.42 |
| | Asn | 341 | OD1 | 2.36 | 2.38 | 2.38 | 2.36 | 2.40 | 2.41 | 2.46 | 2.38 | 2.40 | 2.38 | 2.40 | 2.38 | 2.43 | 2.41 | 2.45 |
| | Asp | 342 | OD1 | 2.34 | 2.33 | 2.33 | 2.35 | 2.34 | 2.30 | 2.35 | 2.32 | 2.28 | 2.35 | 2.30 | 2.33 | 2.44 | 2.30 | 2.31 |
| | Asp | 342 | O | 2.57 | 2.56 | 2.58 | 2.62 | 2.54 | 2.60 | 2.62 | 2.59 | 2.59 | 2.55 | 2.54 | 2.53 | 2.59 | 2.56 | 2.57 |
